# Somatostatin interneurons select dorsomedial striatal representations of the initial learning phase

**DOI:** 10.1101/2024.09.27.611194

**Authors:** S Rotariu, G Zalcman, N Badreddine, F Appaix, S Sarno, I Bureau, E Fino

## Abstract

The dorsomedial striatum (DMS) is an associative node involved in the adaptation of ongoing actions to the environmental context and in the initial formation of motor sequences. In early associative or motor learning phases, DMS activity shows a global decrease in neuron firing, eventually giving rise to a select group of active cells, whose number is correlated with animal performance. Unveiling how those representation emerge from DMS circuits is crucial for understanding plasticity mechanisms of early adjustments to learning a task. Here, we hypothesized that inhibitory microcircuits formed by local interneurons are responsible for the genesis of early DMS representation and associated task performance. Despite the low density of somatostatin (SOM)-positive cells, we observed that selective manipulation of SOM cells disrupted reorganization of DMS activity and modulated initial phases of learning in two behavioral contexts. This effect was cell-specific as manipulation of parvalbumin-positive interneurons had no significant effect. Finally, we identified the high plasticity of SOM innervation in the DMS as a key modulator of the SPN excitability and firing activity. Hence, SOM interneurons set the pace of early learning by actively controlling the remapping of DMS network activity.

## INTRODUCTION

The dorsal striatum, historically considered a critical brain structure for normal motor control, is also acknowledged for its influence on integrative functions and motivational aspects of behavior (Ito and Doya, 2011; Kravitz and Kreitzer, 2012; Liljeholm and O’Doherty, 2012). This progress in understanding has come as functional subdivisions within the striatum have highlighted putative specific contributions to various aspects of a behavior. The dorsomedial part of the striatum (DMS) is involved in the simultaneous integration of cognitive, proprioceptive or sensory information and motor control (Hintiryan et al., 2016; Hunnicutt et al., 2016), constituting an associative node. By this means, it performs multimodal integration (Nagy et al., 2005; Reig and Silberberg, 2014), strongly congruent with locomotion and motor control (Barbera et al., 2016; Fobbs et al., 2020; Kravitz et al., 2010; Nagy et al., 2018; Tecuapetla et al., 2016) and favor early stages of motor sequences formation (Kupferschmidt et al., 2017; Yin et al., 2009). The associative striatum thus appears central in integrative processes essential for developing and adapting motor programs; but how it is expressed within DMS activity patterns is still unclear. During initial phases of associative learning or motor learning, DMS neural activity displays an overall transitory decrease (Badreddine et al., 2022; Cataldi et al., 2022; Vandaele et al., 2019). In addition, in our previous findings on motor learning, we showed that such early DMS activity decrease unveils the emergence of a sparse set of highly active (HA) cells, whose number is directly linked to performance, thus forming representations of the initial training phase within the DMS (Badreddine et al., 2022). These representations could correspond to the adjustment of action responses during the early stage of performance improvement. To explore this, we searched for the precise neural mechanisms that allow the emergence of these representations from DMS activity and evaluated their impact on mouse behavior.

We propose the recruitment of local inhibitory microcircuits, potently regulating the striatal projection neurons (SPNs), as a central mechanism. Local GABAergic interneurons are sparse and form heterogeneous microcircuits of mixed cell types with different distribution and functional output (Assous and Tepper, 2019; Silberberg and Bolam, 2015; Tepper et al., 2018). We focused on two subtypes, somatostatin (SOM) and parvalbumin (PV) interneurons. In the DMS, PV cells have a lower density than in the dorsolateral striatum (DLS) (Bernacer et al., 2012; Fino et al., 2018; Gerfen et al., 1985). On the contrary, while having similar density in DMS and DLS, SOM cells are particularly excitable in DMS, with a greater control over DMS SPNs (Fino et al., 2018). Their spontaneous tonic activity, reported both *ex vivo* and *in vivo* (Beatty et al., 2012; Sharott et al., 2012), is also higher in DMS (Fino et al., 2018). Rare functional studies on SOM interneurons show that a complete ablation of DLS SOM cells does not induce strong motor impairment (Gazan et al., 2020), while activity of DMS SOM cells has been linked to reward-associated instrumental learning (Holly et al., 2019). We thus asked whether SOM microcircuits were able to select DMS neuronal representation of early training and thereby to modulate the associated performance in motor learning or spontaneous exploration of an open field.

We report that selective chemogenetic manipulation of DMS SOM cells precluded the remapping of DMS activity associated with the early training. At the behavioral level, it altered the initial rate of acquisition of the rotarod task and the initial exploration strategy of an open field. Our results suggest that in the two different behavioral contexts, activation of SOM cells leads to an optimized strategy that allows a faster acquisition of the tasks. This control was specific to SOM cells, no behavioral or activity changes were observed as a result of DMS-PV cell manipulation. Finally, we identified several mechanisms in the connectivity plasticity during early training, such as a more efficient feed-forward inhibition from cingulate inputs and an increased SOM-SPN connections which led to a modulation of DMS SPN excitability. Taken together, the data suggest that DMS-SOM microcircuits select task-relevant DMS populations and control the associated behavior.

## RESULTS

### SOM cells silencing prevents the formation of early training DMS representation

We first tested whether SOM cell activity controlled DMS functional remapping and the emergence of neuronal representations of early rotarod training (10 Trials, 1 Session). We used the inhibitory DREADDs system in combination with *ex vivo* calcium imaging: SOM-cre mice were co-injected with AAV-GCaMP6f and AAV-DIO-mCherry or AAV-DIO-hM4Di-mCherry and trained for one rotarod session (10 trials) (Fig. 1A). We built functional maps of evoked DMS activity, by combining the spatial distribution of SPNs within the DMS field of view and the amplitude of the cortically-evoked response in each SPN (Fig. 1B). We compared the network activity of naive mice (mCherry naïve), early-trained control mice (mCherry early-trained) and early-trained mice in which SOM cells were inhibited during each trial of the training (hM4Di early-trained) (Fig. 1B).

**Figure 1.**
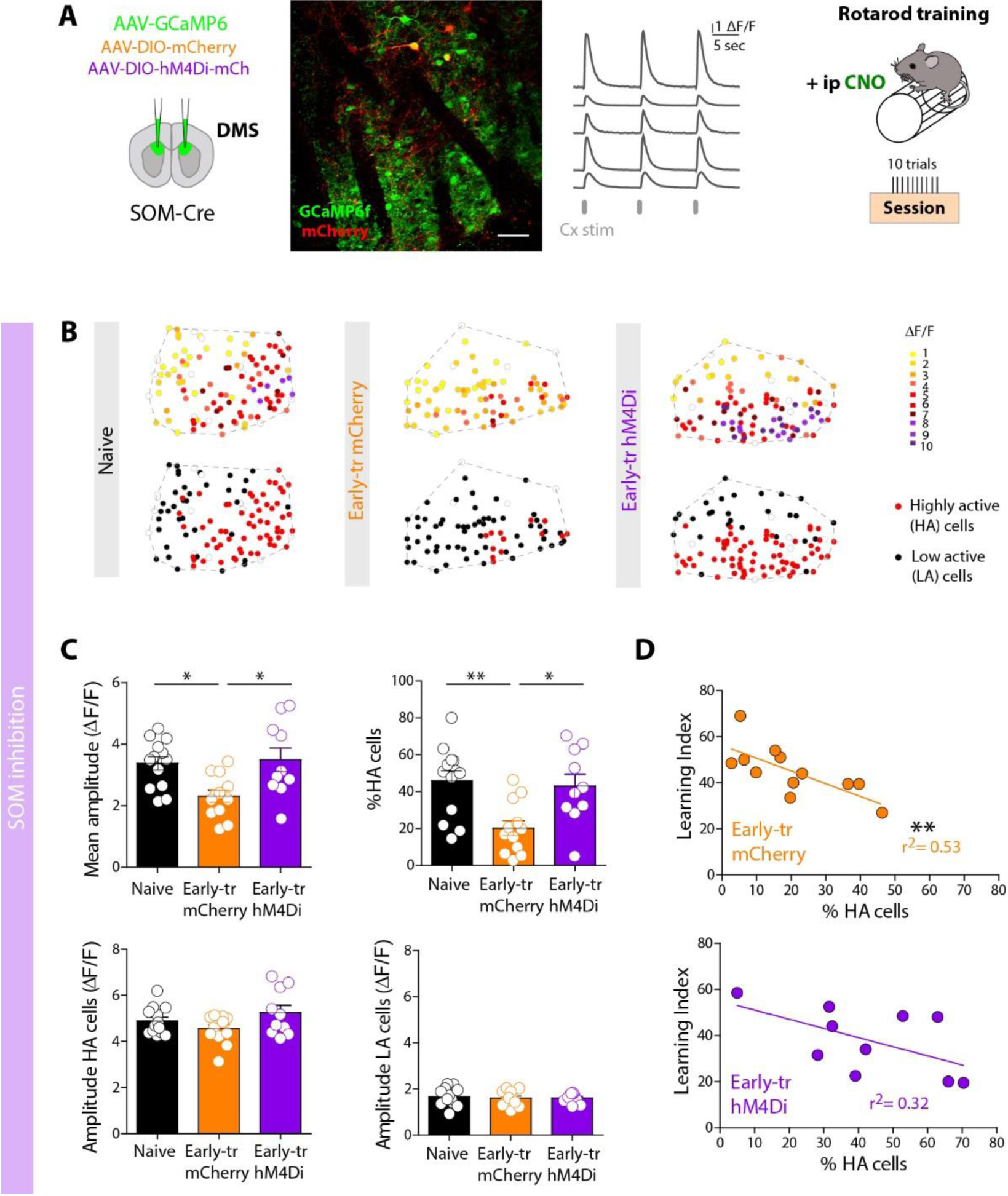
SOM interneurons control the DMS remapping associated with early motor training. A, Experimental conditions. Injections of GCaMP6f in SOM-cre mice with either floxed mCherry (control group) or floxed hM4Di (SOM silencing group). CNO triggers hM4Di mediated inhibition of SOM cells during rotarod training (1 session, 10 trials). The following day, *ex vivo* calcium imaging is performed on three different groups: naive, early-trained or trained with SOM silencing. Two-photon image of GCaMP6f-expressing DMS striatal cells and SOM interneurons labelled with mCherry (scale bar: 50 µm). B, Representative functional maps of the DMS evoked activity in the 3 groups. Top are maps with color-coded amplitude (ΔF/F) and at the bottom, highly active (HA) cells are represented in red and low-active (LA) cells in black. C, For the three different groups: naive (not trained, black, n=5 mice), early-trained (early-tr mCherry, orange, n=11 mice) or trained with SOM silencing (early-tr hM4Di, purple, n=10 mice). Mean amplitudes of all SPNs responses: naïve, 3.36±0.22, n=1143 cells, n=13 slices, n=5 mice, mCherry early-tr, 2.31±0.20, n=1054 cells, n=12 slices, n=11 mice and hM4Di early-tr, 3.49±0.39 ΔF/F, n= 1022 cells, n= 10 slices, n= 10 mice; p= 0.0029, One-way Anova, with Tukey post-hoc. Percentage of HA cells, (Naïve, 45.93±5.31 %, n= 13 slices, mCherry early-tr, 20.22±4.08 %, n= 12 slices, hM4Di early-tr, 43.06±6.41 %, n= 10 slices, p= 0.0023, one-way Anova, Tukey post-hoc). Amplitude of HA cells: Naïve, 5.03±0.25 ΔF/F, mCherry early-tr, 4.56±0.18, hM4Di early-tr, 5.25±0.32, p=0.149, one-way Anova Amplitude of LA cells : Naïve, 1.66±0.11 ΔF/F, mCherry early-tr, 1.59±0.09, hM4Di early-tr, 1.61±0.07, p=0.877, One-way Anova D, Correlation between the percentage of HA cells and the learning index (LI) of the mice after early training (orange) or early training coupled to SOM silencing (purple). The correlation is significant for the early-trained group (top, r^2^= 0,53, p=0,0076, Pearson correlation) but not for the SOM silencing group (bottom, r^2^= 0,32, p=0,0903).

Overall DMS activity was assessed by the mean amplitude of cortically-evoked calcium responses in all SPNs. The mean amplitude was significantly reduced in the early-trained animals, compared to naïve mice, as previously described (Badreddine et al., 2022). Notably, SOM cell silencing had a marked effect since hM4Di early-trained animals had significantly higher amplitude compared to early-trained group, which was similar to the level of naïve animals (Fig. 1C). Out of the global decreased activity, we confirmed that early training neuronal representations where constituted by few particularly highly active cells (HA cells) (Fig. 1C). Silencing SOM cells during early training (hM4Di early-tr) resulted in a significantly higher percentage of HA cells compared to the early-trained group, this percentage was equivalent to naïve animals (Fig. 1C). There was no difference in the amplitude of HA cells, or low active cells (LA cells). Thus, silencing DMS-SOM cells prevented the early DMS remapping. Consistent with the emergence of HA representation of the early phase of motor learning, the mCherry group showed a significant correlation between the percentage of HA cells and the learning index of each animal (Fig. 1D, top). On the contrary, there was no significant correlation in the hM4Di group (Fig. 1D, bottom). To validate an impact of SOM cell activity on neighboring SPNs, we showed that the activation of SOM cells (with excitatory DREADDs) in naïve animals led to a decrease of DMS overall activity level, leading to sparser active SPNs (Supplementary Fig. 1). The same experiments with the activation of PV interneurons had no significant effect (Supplementary Fig 1).

### DMS-SOM cells activity bidirectionally modulate the initial rate of motor learning

Our next aim was to assess whether there was a direct causal relationship between the control of DMS remapping by SOM cells and behavioral performance. Over a full course of rotarod training (5 consecutive days, one session a day, 10 trials per session), we tested the effect of DREADDs-induced manipulation of DMS-SOM microcircuits on the evolution of mouse performance (Fig. 2 and Supplementary Fig. 2). We compared one control group (SOM-cre mice injected with AAV-DIO-mCherry) and a test group in which DMS-SOM cells were silenced during each session (group injected with AAV-DIO-hM4Di-mCherry). Both groups steadily improved their rotarod performance over the 5 days of training. However, silencing SOM activity significantly decreased the initial learning rate as shown by the difference in the latency-to-fall curves (Fig. 2B). In addition, we used quantitative measure with one-phase association fit and observed a significant change in the slope (Fig. 2B-C and Methods). The hM4Di group showed a lower level of performance on Day 1 (Fig. 2D) and reached the level of performance of the control mCherry group on Day 5 (Fig. 2E). This suggests that DMS-SOM activation is required during the early phase of a novel motor learning task. The effect appears to be SOM-specific, given no significant changes on the learning rate were observed when experiments were repeated with PV interneuron silencing (Supplementary Fig. 3).

**Figure 2.**
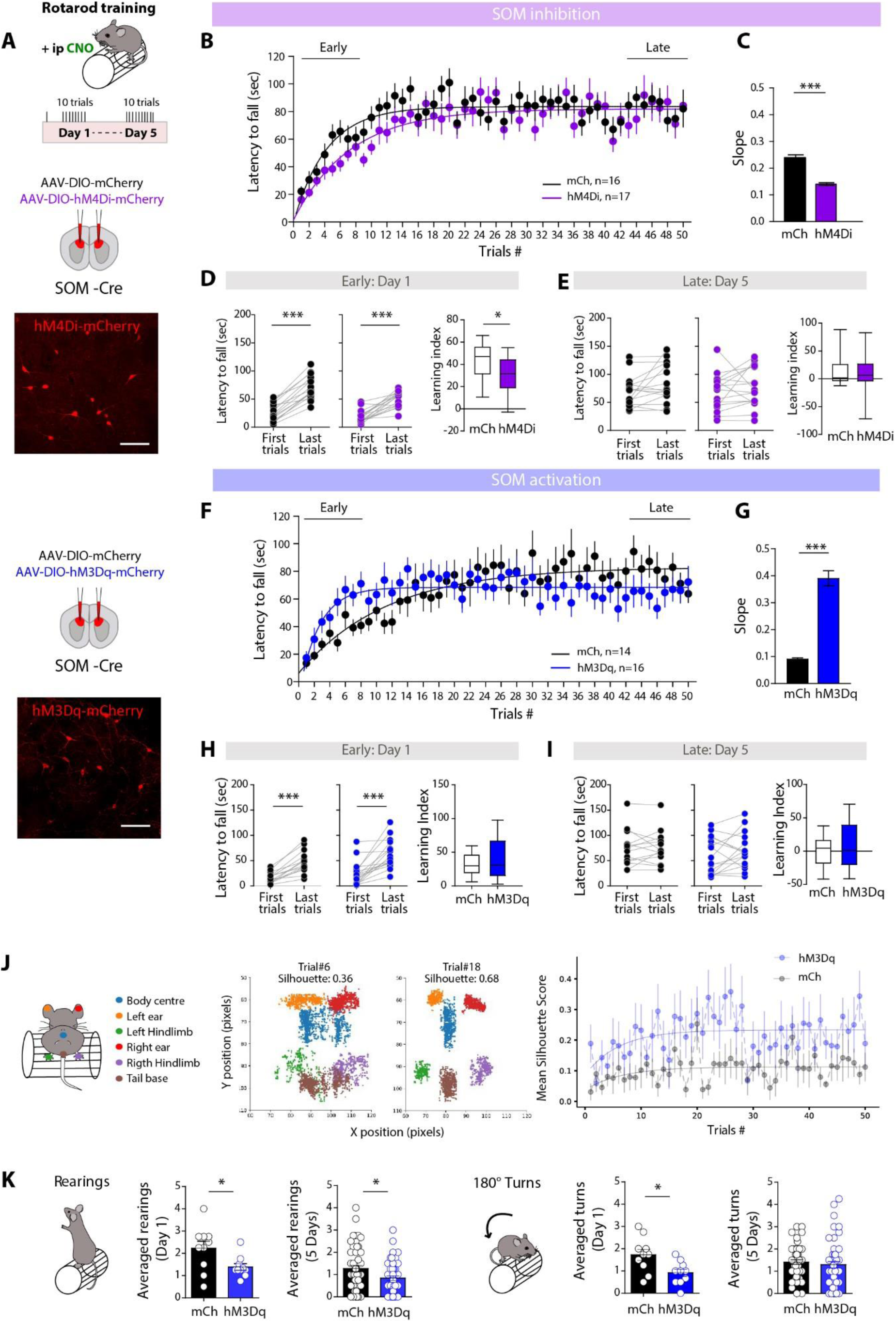
DMS-SOM cells bidirectionally modulate the rate of early motor learning. A, Experimental conditions, SOM-cre mice are injected with either floxed mCherry (control group) or floxed hM4Di (SOM silencing group) or floxed hM3Dq (SOM activating group) during rotarod training (5 days, 1 session with 10 trials per day). Confocal images of SOM cells expressing hM4Di or hM3Dq (scale bars: 50 µm). B, Learning curves and fits employing a one-phase association model represented by the equation y = y0 + (ylim - y0) * (1 - exp(-k * t)) for the control mice (mCherry, black, n=16 mice) or SOM silenced mice (hM4Di, purple, n=17 mice) throughout the 5 training sessions (50 trials). C, Slopes of the one-phase association fits for both groups. mCherry: 0.24±0.04, n=16, hM4Di: 0.14±0.02, n=17, p<0.0001, unpaired t-test. D, In early (Day 1) training, significant improvement in the mice performance between the first and last trials for both groups (mCherry, p< 0.0001, hM4Di, p< 0.0001, paired t-test) but the learning index was significantly lower for the hM4Di group (p= 0.0315, unpaired t-test). E, Both groups reach similar performance by the late phase of the training (Day 5), mCherry, p=0.0588, hM4Di, p= 0.2929, paired t-test. F, Learning curves and one-phase association fits of the control mice (mCherry, black, n=14 mice) or SOM activated mice (hM3Dq, blue, n=16 mice) throughout the 5 training sessions (50 trials). G, Slopes of the one-phase association fits for both groups: (mCherry: 0.09±0.02, n=14 mice, hM3Dq: 0.39±0.11, n=16 mice, p<0.0001, unpaired t-test). H, In early (Day 1) training, significant improvement in the mice performance between the first and last trials for both groups (mCherry, p< 0.0001, hM3Dq, p= 0.0001, paired t-test) with higher performance in hM3Dq group. Learning index was not significantly different (p= 0.4079, unpaired t-test). I, Both groups then reach similar performance by the late phase of the training (Day 5), mCherry, p=0.7309, hM3Dq, p= 0.2492, paired t-test. J, The positions of six body parts were monitored in videos across 50 trials of rotarod training. Representative scatterplots illustrating the XY plane tracking of the body parts for the same mouse across different trials, here trial#6 (left, silhouette value= 0.36) and trial#18 (right, silhouette value= 0.68) of a hM3Dq mouse. Below, averaged silhouette clustering curves were generated for each group (n= 9 mCherry mice and n= 8 hM3Dq mice), and fitted employing a one-phase association model represented by the equation y = y0 + (ylim - y0) * (1 - exp(-k * t)). Permutation test was employed to assess the differences in the three fitted parameters between the two groups: y0: p=0.291, ylim: p=0.055, k: p=0.480 (See Methods). K, Numbers of rearings and turns the mice displayed on the rotarod. In the first session, hM3Dq mice displayed less rearings (hM3Dq, 1.38±0.15, and mCherry, 2.23±0.32, n=10 trials for each group, n=4 mice per group, p=0.048, t-test) and fewer turns (hM3Dq, 0.92±0.16, and mCherry, 1.73±0.25, n=10 trials for each group, n=4 mice per group, p=0.021, t-test). During the full 5 sessions, rearings (for hM3Dq, 0.84±0.10 and for mCherry, 1.27±0.14, n=50 trials for each group, n=4 mice per group, p=0.036, t-test) and turns (hM3Dq, 1.30±0.15 and for 1.41±0.11, n=50 trials for each group, n=4 mice per group, p=0.166, t-test).

To validate these observations, we tested the opposite strategy, i.e. activation of SOM cells during training. The control group was injected with AAV-DIO-mCherry and the test group with AAV-DIO-hM3Dq-mCherry (excitatory DREADDs) (Supplementary Fig. 2). Following the same training protocol, we observed that the hM3Dq group had a significantly improved performance during the first day of training, reaching their performance plateau faster than the control mCherry group (Fig. 2F-I). The slopes of one-association fit of the latency-to-fall curves showed a significant difference. Again, both reached a similar level of performance on Day 5. Replication of these experiments while activating PV interneurons did not significantly affect the learning rate, confirming a SOM-specific role in early motor learning performance (Supplementary Fig. 3).

In the initial phases of the rotarod task, animals must adapt their body positions, to develop appropriate movements and postures that will prevent them from falling. To further analyze variations in behavioral performance, we investigated whether the greater performance of the hM3Dq group correlated with a better positioning of the mouse on the rod. We tracked the positions of six key body parts (2 ears, body center, tail base, and 2 limbs) using DeepLabCut (Lauer et al., 2022). We assessed the degree of organization by analyzing scatter plots of these 6 body parts in the XY plane and calculating silhouette scores, which measure how well body parts split in different clusters in the XY plane (see Methods). This score gives therefore a measure of the whole body positioning and stability on the rod, and not the dynamics of a single body part. Silhouette scores of this scattering increased with learning before reaching a plateau, suggesting that mice increased their stability by confining the movements of each body part in distinct domains of the rod (Fig. 2J, middle panels). The silhouette learning curves (Fig 2J, right panel) reached a higher plateau for hM3Dq mice compared to mCherry mice. The hM3Dq mice thus tended to show more optimized positioning over the course of training than the control mice. To further evaluate the optimization in performing the task, we investigated other side behaviors that were not directly related to the walking on the rod. We quantified the number of rearings and turns (180° turns on the rod), on Day 1 and 5. We observed that SOM activation significantly decreased the number of rearings on Day 1 and over the course of the 5 days (1.27±0.14 for mCherry, 0.84±0.10, n=50 trials for each group, n=4 mice per group, p=0.036, t-test) (Fig. 2K). It also significantly decreased the number of turns on Day 1, but did not change it over the course of 5 days. Conversely, SOM silencing had the opposite effect, a significant increase in the number of rearings compared to control animals on 5 Days (Supplementary Fig. 4). Overall, these data suggested that SOM activation could improve performance on the task by making the mice well positioned and more focused, from the first trials.

### SOM cells activation shortens the discovery phase during an open field task

To evaluate the impact of SOM interneurons in a different context, we compared the spontaneous behavior of hM3Dq and mCherry mice while exploring an open field (OF), i.e. when first introduced to a novel environment (Fig. 3A). Mice from both groups spent a similar amount of time in the center suggesting that the activation of SOM cells had no effect on the animals’ level of anxiety (Fig. 3B). However, hM3Dq mice were ∼ 15 % slower than control mice during the first few minutes of the session (Fig. 3C). This difference was transient because the mean speed measured over the entire session was similar in the two groups of mice (p > 0.05; Fig. 3C), as well as total travelled distance (Supplementary Fig. 5), showing that none of the group displayed hyper- (or hypo-) locomotion. This finding suggested an effect of the SOM cell activation on the way the animal explored the open field during the first few minutes of discovery of the arena. To investigate this further, we considered two speed ranges (corresponding to the two peaks of speed distribution in Fig. 3C and associated with two behavioral states (Sheets et al., 2013)). We observed that hM3Dq mice were more frequently in the slow mode early in the session, especially when at the center of the OF (Fig. 3D). The slow and fast modes corresponded to two different types of trajectories during exploration; in fast mode, mice ran in straight lines whereas in slow mode their trajectories were more tortuous (Fig. 3E). Thus, the increased prevalence of the slow mode suggested a different spatial organization of the hM3Dq mice displacements. To test this, we analyzed for each mouse the sequence of its displacement by counting for each visited zone the number of transitions that were made for the animal to return (Fig. 3F-G). If the number of transitions was low, it would mean that the animal focuses on some local regions, and, on the contrary, if the number of transitions was high, it would indicate wide displacements across the OF. For the mCherry mice, this number was high at the beginning and decreased to stabilize after ∼ 4 minutes. In contrast, for the hM3Dq mice, the number of transitions was already low at the beginning, close to the number found later in the session (Fig. 3F-G). These results thus suggest that the hM3Dq mice used different strategies to explore the open field, at a slower pace and with more local inspection of some OF zones, while control mice start with a wide exploration of the OF before restricting the zones of interest. Importantly, we observed no difference in the exploration of the OF when DMS-PV cells were activated or DMS-SOM cells were inhibited (Supplementary Fig. 5). Taken together, these results suggest that SOM activation modified the exploratory behavior of hM3Dq mice while discovering a novel environment bringing it to a state that was normally observed later in the session.

**Figure 3.**
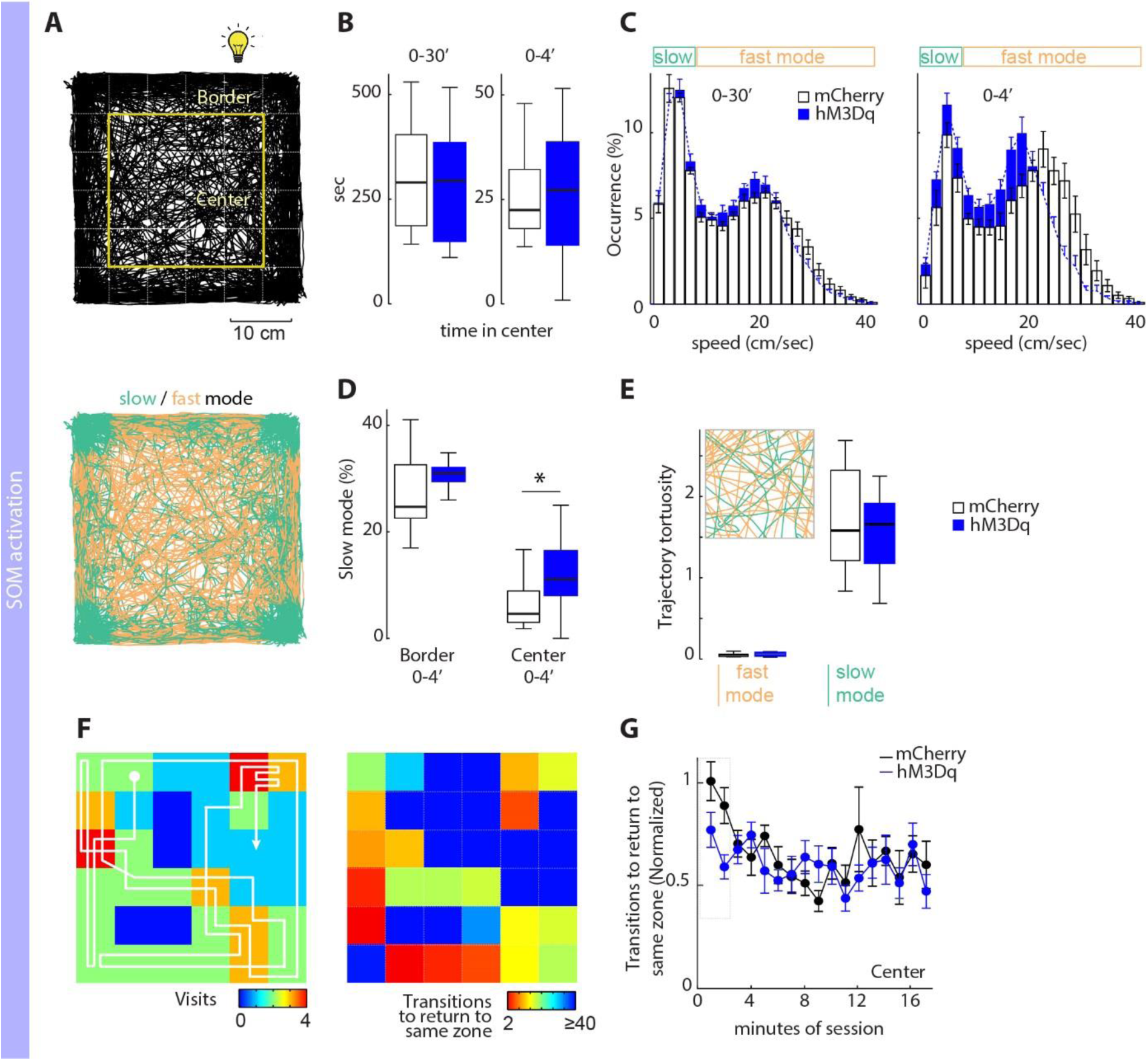
SOM cells impact the mouse behavior at the beginning of an open-field exploration. A, Superposed trajectories of 11 hM3Dq mice (top) acquired during the first 4 min of the open-field (OF) exploration with limits of center and border of the OF. Dashed lines separate the arena in 36 zones used for the analysis. Below, same trajectories color-coded for the mouse speed. B, Time spent in the center of the OF during the entire session (left) and the first 4 minutes (right) for mCherry (white) and hM3Dq (blue) mice. C, Histograms of the mouse speed during the entire session (left) and the first 4 minutes (right) suggest two speed modes, below 8 cm/sec (slow) and above 8 cm/sec (fast). The hM3Dq mice (blue) are slower in the first 4 minutes of exploration: mCherry, 17.2 ± 0.8 cm/sec, n= 11 mice; hM3Dq, 14.3 ± 0.4 cm/sec, n=13 mice; p = 0.0205. This is significant both at the border (mCherry, 15.8 ± 0.8 cm/sec, n= 11 mice; hM3Dq, 13.4 ± 0.4 cm/sec, n=13 mice; p = 0.043) and in the center of the OF (mCherry, 20.8 ± 0.9 cm/sec; hM3Dq, 17.1 ± 0.8 cm/sec; p = 0.015). D, Fraction of zones visited in slow-mode during the first 4 minutes is higher in the hM3Dq mice in the center only (p = 0.049). E, The tortuosity of trajectories is higher in slow mode, in both mCherry and hM3Dq. Inset, close up view (8 x 8 cm) of trajectories at the center shown in A. F, Left, trajectory of a single hM3Dq mouse (white line) and the number of visits of each zone during the first minute of exploration, reported in a color map. Right, for the same mouse, color map of the number of transitions made before returning to a given zone. Warm colors designate temporary «hot-spots», zones where the animal returned with fewer transitions during this period. G, Number of transitions to return to an already visited zone normalized for the total number of transitions made at each time bin. Values < 1 indicate a preference for recently visited zones and values > 1 indicate avoidance of these zones. Dashed box indicates significant difference (Anova, p = 0.006).

### SOM interneurons efficiently modulate SPN excitability in DMS

Calcium imaging data showed that SOM cells were able to select SPN subpopulations (HA cells) to form early learning representations. We thus investigated by which mechanisms they could increase the signal-to-noise ratio and control the patterning of DMS activity. One hypothesis is that they would potently modulate the excitability level of DMS-SPNs. Comparing the intrinsic properties between HA and LA cells after the training showed no difference (Badreddine et al., 2022). This suggested that, if true, the modulation of SPN excitability might be triggered during the training itself. We thus compared the intrinsic membrane and firing properties of DMS SPNs of before (in naïve mice) and after early-training, using patch-clamp recordings in acute brain slices (Fig. 4A-C). The recorded cells were mainly LA cells (in early-tr condition) as they represent 80% of the SPNs. We observed that early-training by itself led to a decrease in SPN excitability. Indeed, SPNs in early-trained group had a lower input resistance and a higher rheobase (Fig. 4C). There was no significant change in the AP threshold. Although there was no change in the spiking frequency in response to a given current injection, there was a rightward shift in the current/spiking frequency relationship in early-trained mice (Fig. 4B). Thus, early training reduced DMS-SPN excitability.

**Figure 4.**
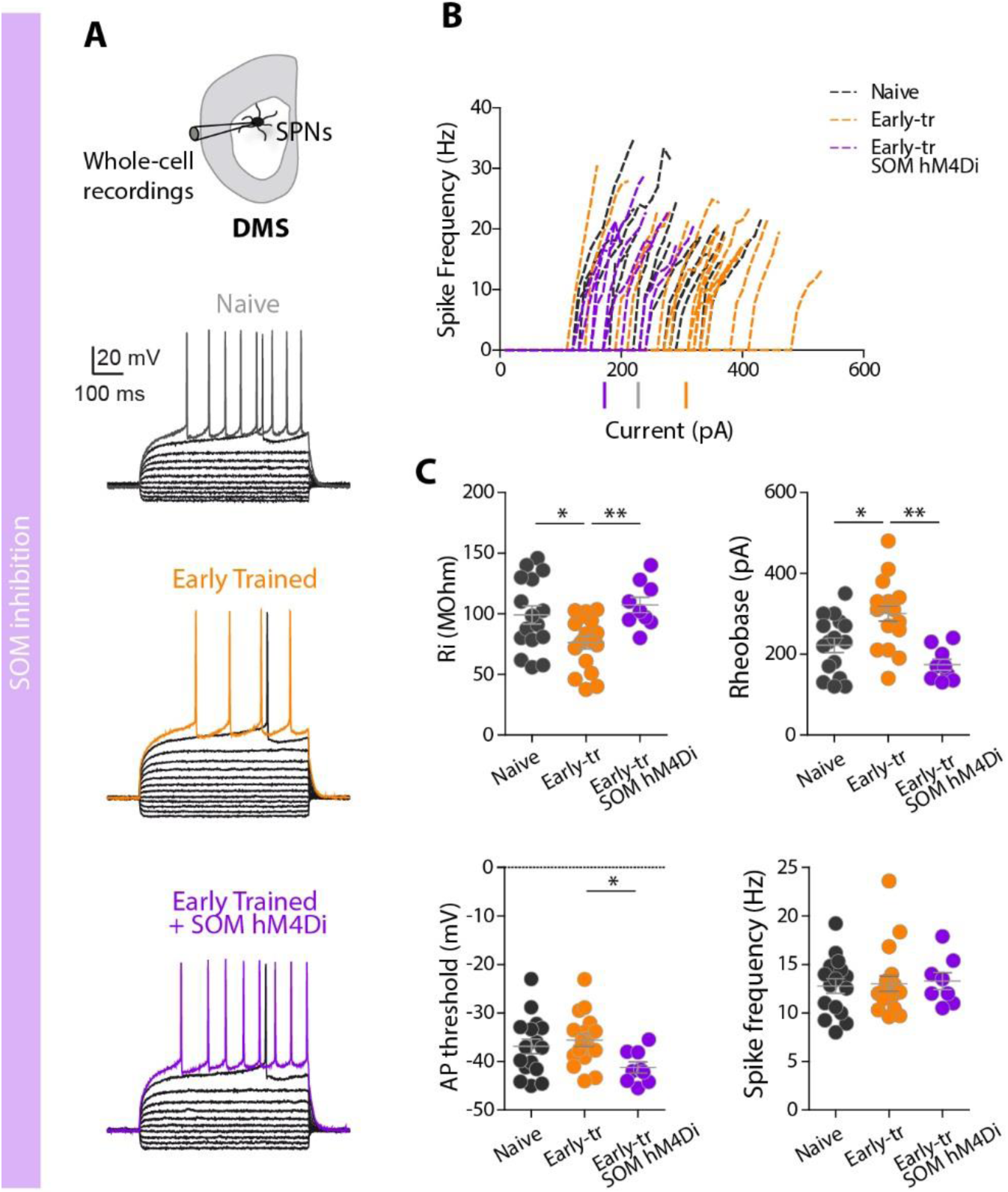
DMS-SOM cells are involved in training-induced decrease in SPN excitability. A, Patch-clamp recordings of SPN in naïve (dark grey), early-trained (early-tr, orange) or early-trained with SOM silencing (early-tr SOM hM4Di, purple) were performed. Representative I/V curves of SPNs in the three conditions. B, SPN spike frequency in relation to the injected increasing current in the three experimental groups. C, SPN intrinsic properties in the different conditions (One-way Anova). Input resistance (Ri): Naïve, 98.94±7.50 MΩ, n=16 SPNs, n=8 mice, Early-tr, 76.14±5.07 MΩ, n=18 SPNs, n=7 mice, Early-tr SOM hM4Di, 107.2±6.3 MΩ, n=9 SPNs, n=4 mice, p=0.005. Rheobase, Naïve, 221.9±18.0 pA, Early, 300.6±19.2 pA, Early-tr SOM hM4Di, 173.9±13.6 pA, p=0.0002. AP threshold: Naïve, −36.9±1.5 mV, Early, −35.6±1.3 mV, p=0.5157, Early-tr SOM hM4Di, −41.2±1.1 mV, p=0.0436. Spike frequency: Naïve, 12.8±0.8 Hz, Early-tr, 13.0±0.8 Hz, Early-tr SOM hM4Di, 13.3±0.87 Hz, p=0.9291.

We next questioned the implication of SOM cells in this decrease in SPN excitability. Similar SPN path-clamp recordings were made in mice early-trained while silencing SOM cells (hM4Di, inhibitory DREADDs) (Fig. 4A-C). We found that SPNs in hM4Di early-trained mice had a higher excitability compared to early-trained animals as evidenced by a higher input resistance and a lower rheobase. Spiking frequency was unchanged but there was a leftward shift in the current/spiking frequency relationship in hM4Di early-trained mice, overlapping with naïve mice (Fig. 4B). Overall, hM4Di early-trained mice exhibited SPN intrinsic properties that were not significantly different from those of naïve mice. Collectively, these results suggest that SOM cells could set up the gain of DMS network activity by efficiently modulating the SPN excitability.

### Training shortens SOM cells latency recruitment by cingulate inputs

We next investigated whether modulation of DMS-SPN excitability was mediated by an enhanced feed-forward inhibition. We asked whether early training would change the dynamics of SOM cells or SPN recruitment by cingulate inputs. To do so, we compared cortically-evoked calcium responses in SPNs and in mCherry-labelled SOM interneurons from naïve and early-trained mice (Fig. 5 A-B). We evaluated whether the proportion of recruited cells, the calcium-evoked response amplitudes and their latency times were affected by training. The vast majority of SOM cells and SPNs were activated by the cortical stimulation in both training conditions (> 90%, Fig. 5 C-D). As described in the first part of the Results section, SPNs showed a significant decrease in the amplitude of their responses between naïve and early trained mice (Fig. 5D). In contrast, SOM cells had similar amplitude calcium responses in naïve and early-trained mice, indicating stable responses. It should be noted that the small amplitude of responses from SOM cells indicates that they were part of the LA population (below the threshold of HA level), so likely out of the HA representations.

**Figure 5.**
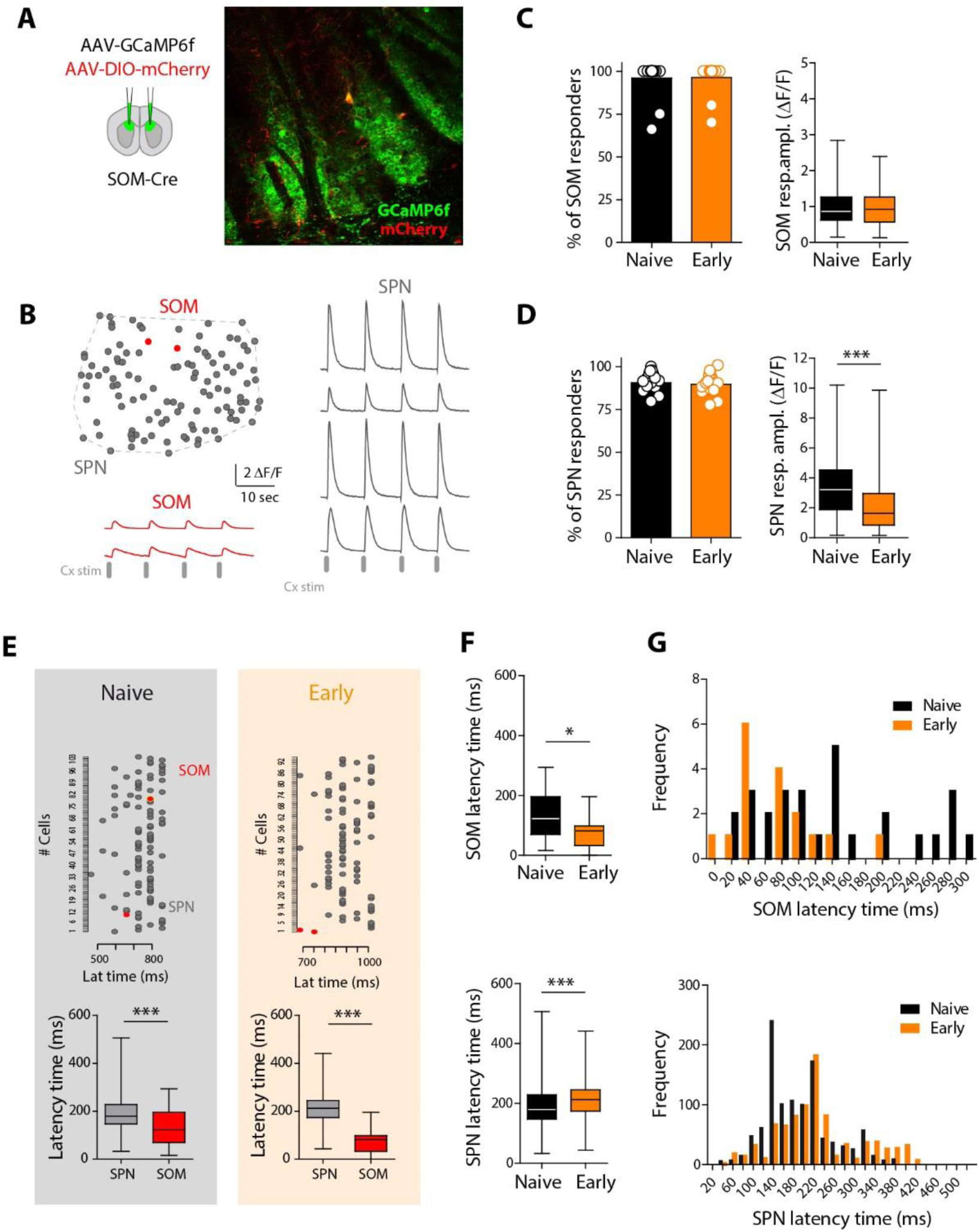
Training oppositely changes the SOM and SPN cortically-evoked calcium responses. A, Experimental conditions: SOM-cre mice are injected with GCaMP6f and with floxed mCherry to identify SOM cells in the recorded FOV. B, Schematic of a FOV with SOM cells (red) and SPNs (grey) and representative cortically-evoked calcium responses. C-D, Percentage of SPN (C) or SOM cells (D) responding to the cortical stimulation in naïve or early-trained (early) mice and mean amplitude of the calcium responses. For both cell types, the percentage of responding was similar between naïve and early-trained mice. For SOM cells, in naïve mice, 96.07±2.72%, n=30 SOM cells from n= 15 slices from n= 5 mice and in early mice, 96.4±2.5%, p=0.971, n=31 SOM cells from n=19 slices from n= 15 mice. For SPNs, in naïve, 91.2±1.4%, n=1143 SPNs from n= 13 slices from n= 6 mice and in early, 93.2±1.1%, n=1503 SPNs from n= 12 slices from n= 11 mice, p=0.278). The mean amplitudes of SPNs evoked responses was significantly modulated by the training stage (p<0.0001) while no difference for SOM cells were shown (p=0.991). E, Raster plots of calcium-evoked responses latency times in SPNs (grey) and SOM cells (red) in naïve or early conditions. Training changes the latency time of calcium-evoked responses. SPNs have longer latency time in naïve (p< 0.001, Mann-Whitney) and early (p< 0.001) conditions. F-G, Early-training significantly shortens the SOM cells latency time (p= 0.0121, n= 28 SOM cells in naïve and n= 17 SOM cells in early), as also illustrated on the distribution on the right. In SPNs, early-training increases the latency times (p < 0.0001, Mann-Whitney, Naïve, n= 1059 SPNs, n=12 slices, n= 5 mice; Early, n= 840 SPNs, n=10 slices, n= 8 mice).

With respect to the latency of evoked-responses, consistent with feed-forward inhibition, SOM cells were activated with shorter latency times than SPNs in naïve mice. Nevertheless this earlier activation was even more pronounced after early training (Fig. 5E). This was due to a significant shortening of SOM cell latencies (∼half compared to naïve mice) (Fig. 5F-G). Indeed, for SPNs, this latency was on the contrary significantly longer (Fig. 5F-G); although there is a large overlap in the latency times of most SPNs in early and naïve conditions, there was a small subset of SPNs with longer latency times. This subset could correspond to the SPNs that have been inhibited by SOM cells after training. Overall, cingulate afferents activated SOM cells earlier than SPNs, a situation that is further enhanced by learning, suggesting a favored position to mediate efficient feed-forward inhibition.

### Reinforced SOM-SPN connectivity after early training

Finally, to complete SOM-mediated feed-forward inhibition plasticity, we tested whether training could trigger plasticity of local DMS SOM-SPNs connections. Hence, SOM cells might tightly control DMS remapping by exerting a stronger inhibition on surrounding SPNs. To study the SOM-SPN functional connectivity, we used optogenetic activation of SOM cells with an AAV-flox-ChR2-mCherry injection in DMS of SOM-cre mice (Fig. 6A). We performed whole-cell patch-clamp recordings of DMS-SPNs and recorded light-induced currents (LED 473 nm). Early training significantly enhanced the light-evoked currents in SPNs (Fig. 6B-C). SOM-SPNs connections were significantly increased after early training, with a ∼1.5 fold higher amplitude of light-evoked currents in early-trained mice. Next, we investigated how this change in synaptic weight affected their local inhibitory control. To this end, SOM cells were opto-activated (300 ms light pulse) during spike trains elicited in SPNs, to investigate the consequences of an increased SOM-mediated inhibition on the SPN firing as it could occur during early training. Normalized firing frequency (during *vs.* before light pulse) was significantly lower in the early-trained mice compared to naïve (Fig. 6D-E), indicating a stronger inhibition of SPN firing frequency. Taken together, these data suggested that SOM cells have a greater inhibitory synaptic weight on the SPN activity after early training, providing an additional mechanism of control of the early motor learning by the SOM microcircuits in the DMS.

**Figure 6.**
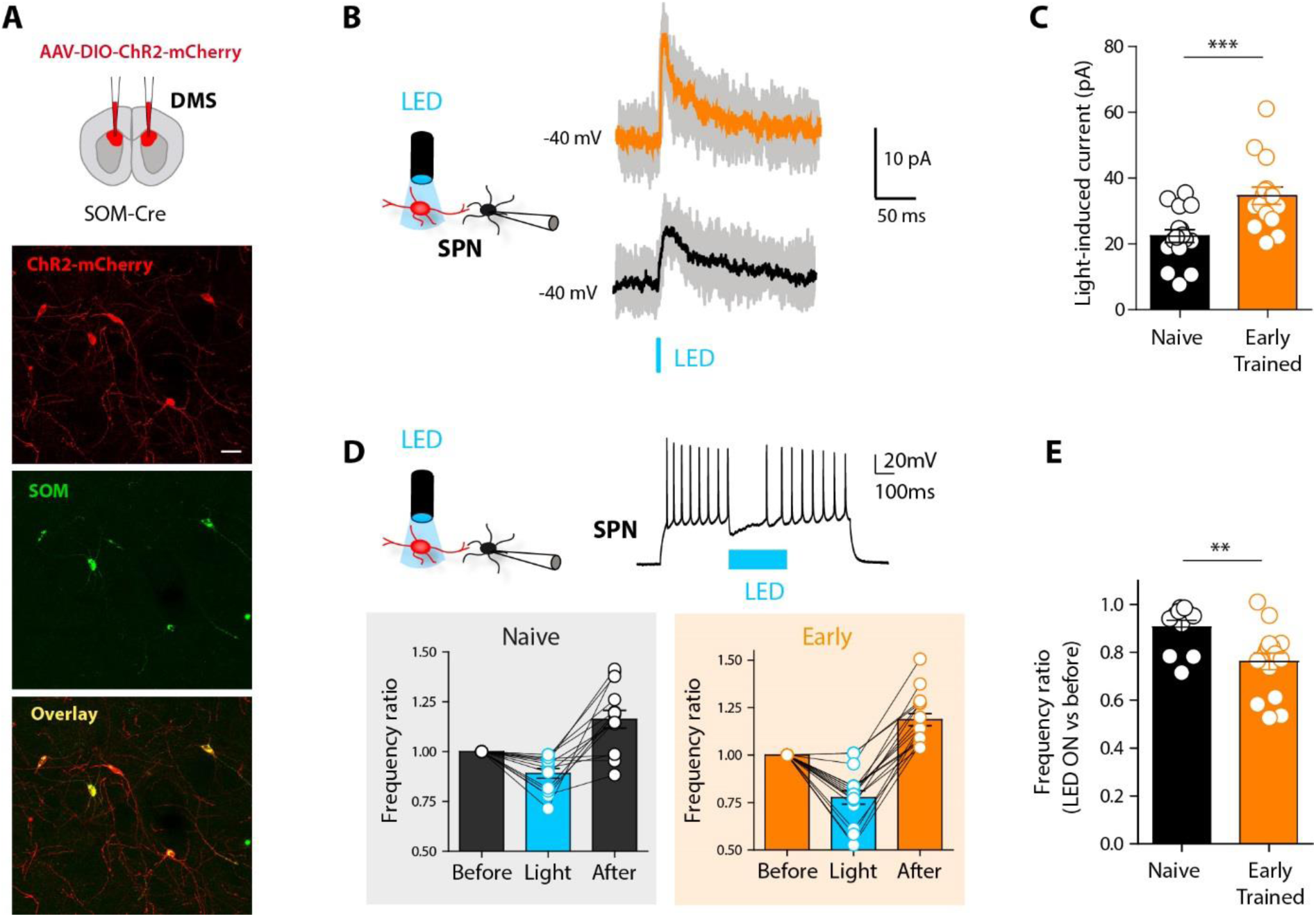
Training increases DMS SOM-SPN functional connectivity. A, Experimental conditions, SOM-cre mice were injected with floxed ChR2-mCherry in DMS. Below, confocal images of SOM cells expressing ChR2-mCherry (scale bar: 20 µm). B, With patch-clamp recordings of SPNs, light-induced current amplitude were monitored in response to short LED pulses (473 nm LED, 5 ms) in naïve (black) or early-trained (orange) mice. The amplitude was significantly higher in early-trained animals: Naïve, 22.42±1.91 pA, n= 17 SPNs, n= 10 mice, Early, 34.68±2.59 pA, n= 16 SPNs, n=7 mice, p= 0.0011, unpaired t-test. D, Depolarizing current steps were applied to the SPN with a long LED pulse (300 ms) in the middle of the spike train. E, The frequency ratio was measured to evaluate the inhibition mediated by the SOM cells to the SPNs. The frequency ratio was lower in early-trained mice, indicating a stronger inhibition: Naïve, 0.90±0.03, n=11 SPNs, n=9 mice, Early, 0.76±0.03, n=16 SPNs, n=7 mice, p=0.0167, Mann-Whitney.

## DISCUSSION

In the present study, we show that DMS-SOM interneurons control initial phases of motor tasks by selecting neuronal representations. SOM-targeted silencing prevents the DMS remapping associated with early training. Interestingly, this control translates to the behavioral performance: activation of DMS-SOM cells shortens (and their inhibition prolongs) the early learning phase of the rotarod training or the open-field initial exploration phase. Whole-cell SPN recordings revealed that these effects are mediated by a decrease of SPN excitability through an enhanced feed-forward inhibition of SOM cells onto SPNs. Notably, such inhibitory control appears to be cell-specific, as similar manipulation of DMS-PV interneurons has no significant effect.

### Control of DMS neuronal ensembles by local SOM interneurons

We describe here how DMS-SOM cells modulate the activity of SPN populations and favor the formation of initial training representations. Such selection by SOM cells is quite striking considering their low density overall within the striatum (Fino et al., 2018; Tepper et al., 2018). Interestingly, some parallel can be made with SOM interneurons in the motor cortex, where they are also required to form and maintain sequential activation of pyramidal cell ensembles associated with motor skill learning (Adler et al., 2019; Yang et al., 2022). The DMS representations reported here are constituted by a subset of sparse, highly active SPNs, whose number is significantly correlated with task performance. Those highly active SPNs represent ∼around 20% of DMS SPNs, coherent with a recent study also reporting a small percentage of SPNs (∼30%) involved in specific aspects of spontaneous or learned motor behavior (Tiroshi, 2023). We previously confirmed with cFos-TRAP labelling of HA cells that they were present *in vivo* and also that these early activity patterns are transient (they are not present in the DMS after longer training, Badreddine et al., 2022). We thus propose that DMS HA cells function could relate to properly indexing or cueing novel environment to the task and that such function relies on SOM cells. Early DMS representations arise from a globally reduced DMS activity, also reported during the early phase of operant conditioning (Vandaele et al., 2019) (Stubbendorff et al., 2019) or learned motor coordination (Cataldi et al., 2022). Interestingly, in our experiments, while DMS mean amplitude is increased after SOM silencing, individual amplitude of highly active or low active cells is not affected. This indicates that the increase in mean amplitude is a consequence of increased percentage of HA cells and not of individual cell amplitude, suggesting an “all or nothing” strategy in which a cell might be chosen (HA cell) or not (LA cell) to be part of the representation by SOM cells inhibition. The global DMS activity decrease could also be favored by corticostriatal synaptic plasticity rules showing a greater occurrence of long-term depression at cingulate-DMS connections (Perez et al., 2022). The LTD amplitude or proportion of SPNs developing LTD would be modulated by interneurons as GABAergic circuits are able to switch plasticity rules through opposite spike-timing dependent plasticity (STDP) learning rules in SPNs and SOM cells (Fino and Venance, 2011; Paille et al., 2013). The effect of SOM cells might rely on a control of SPN spiking through modulation of intrinsic excitability (this report) or efficient feedforward inhibition (Elghaba et al., 2016; Fino et al., 2018; Straub et al., 2016).

Importantly, the control by local inhibitory microcircuits is cell-specific, and not due to any local inhibition since modulation of another interneuron subtype, the PV cells, had no significant effect on network plasticity or behavioral performance. Subtype-specific plasticity of inhibition was also reported in motor cortex after motor learning, with dendrite-targeting SOM interneurons identified as key regulators of learning-associated spine density changes (Chen et al., 2015). With a mild segregation in SPN cellular compartments targeting from striatal PV and SOM interneurons, the SOM-specific modulation might depend on different parameters. Specific electrophysiological, anatomical properties and circuit integration could explain a functional division between interneuron subtypes. Interestingly, SOM cells are particularly abundant in humans (identified as nitrergic neurons) while PV cells density is low in both caudate nucleus and putamen (Bernacer et al., 2012; Lecumberri et al., 2018). This could suggest that the role of SOM/nitrergic neurons in integrative functions might be even more pronounced in humans.

### Training-induced plasticity of striatal SOM cells intrinsic and extrinsic connectivity

Striatal SOM interneurons are integrated into a highly complex circuitry, receiving inputs from GABAergic interneurons (parvalbumin and tyrosine hydroxylase-expressing) and with reciprocal connections with cholinergic interneurons (Assous et al., 2017; Frost Nylen et al., 2021; Lee et al., 2017; Melendez-Zaidi et al., 2019; Szydlowski et al., 2013). Their main targets are the SPNs and although unitary synaptic connections appear weak (Fino et al., 2018; Gittis et al., 2010), optogenetic activation of several SOM cells elicits large inhibitory postsynaptic responses in SPNs (Elghaba et al., 2016; Fino et al., 2018; Straub et al., 2016), showing that SOM cells, as a population, have a significant impact on SPNs. This SOM activity in chorus was also reported in cortical circuits (Safari et al., 2017). We observed that SOM cells are responsible for the decrease in SPN excitability associated with early training, in accordance with more excitable SPNs observed in fully SOM ablated mice (Gazan et al., 2020). Their tonic activity (Beatty et al., 2012; Sharott et al., 2012) and local inhibitory control, greater in DMS, might be responsible for their efficient modulation of SPN intrinsic properties. With long axonal arborizations extending up to 1 mm (Kawaguchi, 1993; Straub et al., 2016), SOM cells can extend their influence over large parts of striatal territories, coherent with the global DMS activity shutdown reported. We also show that SOM-SPNs connections are highly plastic, with enhanced synaptic strength after early training. Plasticity of synaptic inputs modulated by motor skill learning had been previously reported in SPNs (Lopez-Huerta et al., 2021). Changes in synaptic strength is highly reversible and would represent a putative mechanism for transient remapping of the DMS.

SOM interneurons are known to mediate efficient feed-forward inhibition on SPNs (Assous et al., 2017; Assous and Tepper, 2019; Fino et al., 2018; Johansson and Silberberg, 2020). Interestingly, we observed that training efficiently modulate the recruitment of both SPNs and SOM cells and in consequence the efficiency of SOM feed-forward inhibition. Indeed, while early training reduces the average amplitude of SPN calcium transients compared to naive animals, SOM cells are recruited with similar amplitudes and shorter latencies, placing them in a favored position to mediate increased feedforward inhibition. Combined with an increased SOM-SPN connectivity, the plasticity of multiple circuit loci constitute the mechanisms for SOM-mediated DMS remapping.

### Control of initial phases of behavioral tasks

DMS has a role in action-outcome associations (Balleine and O’Doherty, 2010; Morris et al., 2022), which could be associated with the accumulation of evidence and cues from the environment. This idea is supported by a recent study showing DMS activity modulation during a T-maze task, which is dependent on task demands, i.e. with progressive accumulation of cues, versus fixed cues present in the environment (Bolkan et al., 2022). In the two behavioral contexts used in our study, activation of SOM cells leads to an optimized strategy that could mimic a faster accumulation of knowledge and, ultimately, faster initial acquisition of the tasks. Our work thus suggests that SOM cells would be determinant in the selection of DMS-SPN ensembles involved in the discovery phase of the tasks. Nevertheless those tasks do not allow us to distinguish between motor refinement and better integration and adaptation to environmental cues. We did not observe strong effects on overall motor abilities or locomotion in control or SOM-activated mice, but other changes associated with SOM activation led to faster acquisition of the rotarod task and a narrower exploration of the open-field. One could postulate that SOM cells activity would lead to a decrease in behavioral flexibility and/or a transient reduction in exploratory behavior in order to be more focused from the outset. Indeed, the effects of SOM cell activity were significant in the early phase of motor training and first exploration of an open-field, without any significant effect on later stages of the tasks. Such temporally restricted effect was also reported for DLS-PV cells, which influence performance in reward conditioning but only at initial stages as learning decrease the influence of PV on DLS-SPNs (Lee et al., 2017). Within the DMS, SOM (LTS) cells were also described as potent modulators of goal-directed learning (Holly et al., 2019) but, in this context, without time restriction in the learning process.

In spontaneous behavior, a differential contribution of striatal subtypes has been reported; SPNs and PV interneurons (FSIs) would preferentially encode locomotion whereas other interneurons (possibly involving SOM cells) encode environmental identity (Yamin et al., 2013). Accumulating evidence has been linked to cognitive load required to coordinate behavior (Bolkan et al., 2022; Thorn et al., 2010; Yartsev et al., 2018). Bolkan et al suggest that low cognitive load could lead to sparse recruitment of DMS-SPN ensembles whereas higher cognitive load would increase ensemble size recruitment. We cannot conclude from our experiments the contribution of specific subsets of SPNs. While we did not see any strong bias toward active d- or iSPNs in the composition of early DMS representations (Badreddine et al., 2022), it is difficult to draw general conclusions as the balance of activity between dSPNs and iSPNs might depend on the task and its environmental context (Cuevas N., 2024). Further studies dissecting specific behavioral conditions are needed to elucidate the precise contribution of striatal subtypes and neuromodulation to the initial adaptation to a given task, the transition to full engagement, and finally the refinement of the behavior.

## MATERIALS AND METHODS

C57BL6 mice (*Mus musculus*) of 1.5 to 2.5 month-old of both sexes were used and housed in temperature-controlled rooms with standard 12 hours light/dark cycles and food and water were available *ad libitum*. PV-Cre and SOM-Cre lines were used in most of the study (mouse lines #008069 and #013044 from Jackson Laboratory). Every precaution was taken to minimize stress and the number of animals used in each series of experiments. All experiments were performed in accordance with EU guidelines (directive 86/609/EEC) and in accordance with French national institutional animal care guidelines (protocol #29200 and #33185).

## METHOD DETAILS

### AAVs

Adeno-associated viruses (AAVs of serotype 5 and 9) were used to express different genes in striatal cells. AAV5-syn-GCaMP6f-WPRE-SV40, AAV5-hSyn-DIO-hM4D(Gi)-mCherry, AAV5-hSyn-DIO-hM3D(Gq)-mCherry, AAV5-hSyn-DIO-mCherry or AAV9-EF1-dflox-hChR2(H134R)-mCherry were purchased from Addgene (MA, USA).

### Stereotaxic injections

Stereotaxic intracranial injections were used to deliver AAVs in DMS. Mice were anesthetized with isoflurane (5% for induction and procedure at 1-3%) and placed in a stereotaxic frame (Kopf) on a heating pad (37°C). Under aseptic conditions, the skull was exposed and leveled and a craniotomy was made with an electric drill. The viruses (serotype 5/9, ≈ 10^12^ genomic copies per mL) were injected through a pulled glass pipette (pulled with a P-97 model Sutter Instrument Co. pipette puller) using a nanoinjector (World Precision Instruments, Germany). The pulled glass micropipette was slowly lowered into the brain and the injection of the virus was performed at a 100 nL / min rate. A volume of ∼400 nL of the virus was enough to transfect cells from a large proportion of DMS. The DMS injections targeted coordinates AP + 1.2mm, ML 1.2, DV - 2.4. Following injections, the pipette was left in place for 5 min to prevent backflow and was then slowly raised out of the brain. To minimize dehydration during surgery mice received a subcutaneous injection of sterile saline. Postoperatively mice were monitored on a heating pad for 1 h before being returned to their home cage. Mice were then monitored daily for 4-5 days. Behavioral and/or imaging experiments started 15 to 20 days after injection, a period allowing a good expression of AAVs.

### Behavioral training

#### Accelerating rotarod

In the days prior to the training, mice were habituated to the room and to handling. Mice learned to run on a rotating rod (Panlab). For each trial the mouse was placed on the moving rod at the constant speed of 4 rpm. The rotation of the rod was then increasing from 4 to 40 rotations per min over 300 s (Kupferschmidt et al., 2017; Yin et al., 2009). Each trial ended when the mouse fell off the rod or when the 300 s had elapsed. There was a resting period of 300 s between each trial during which mice are put back in the home cage. Mice performed 10 trials per day, for up to 5 days. Early phase corresponds to the 10 trials of the first training day. A subset of animals was video-recorded with a camera (20 fps) controlled by Bonsai software (bonsai-rx.org).

#### Open-field

About 1 week after their last session on the rotarod, animals were injected with CNO (3 mg/kg i.p.) and placed 30 min later in an open-field (40 cm x 40 cm) for 30 min exploratory spontaneous motor activity session while they were video-recorded with Noldus software (Ethovision WT11.5). A small number of trials had missing or disrupted video data and were excluded.

#### *Ex vivo* two-photon imaging and patch-clamp recordings

##### Brain slice preparation

Brain slices preserving DMS and its cortical inputs coming from cingulate cortex were prepared as previously described (Badreddine et al., 2022; Fino et al., 2018). Animals were anesthetized with isoflurane before extraction of the brains. Brain slices (300 μm) were prepared using a vibrating blade microtome (VT1200S, Leica Microsystems, Nussloch, Germany) in a 95 % CO_2_ and 5 % O_2_-bubbled, ice-cold cutting solution containing (in mM) 125 NaCl, 2.5 KCl, 25 glucose, 25 NaHCO_3_, 1.25 NaH_2_PO_4_, 2 CaCl_2_, 1 MgCl_2_, 1 pyruvic acid. Slices were then transferred into the same solution (with 100µM pyruvic acid) at 34°C for one hour and then maintained at room temperature.

##### Two-photon calcium imaging

Genetically-encoded Ca^2+^ indicator GCaMP6f was used for calcium imaging of somas of striatal cells. GCaMP6f was expressed with recombinant AAVs injected in DMS. Two-photon calcium imaging was performed at λ= 940 nm with a TRiMScope II system (LaVision BioTec, Germany) using a resonant scanner, equipped with a 20x/1.0 water-immersion objective (Zeiss) and coupled to a Ti:Sapphire laser (Chameleon Vision II, Coherent, > 3 W, 140 fs pulses, 80MHz repetition rate). The laser power was set at ∼40-50 mW on sample. Fluorescence was detected with a GaAsP detector (Hamamatsu H 7422-40). Scanning mode and image acquisitions were controlled with Imspector software (LaVision BioTec, Germany) (15.3 frames per second for 1024 × 1024 pixels, between 50 to 150 µm underneath the brain slice surface, with no digital zoom). Typical field of view for calcium imaging was 392 x 392 µm. Cortically-evoked activity was recorded in response to electrical stimulations applied with a bipolar electrode (MicroProbes, USA) placed in the layer 5 of the cingulate cortex as previously described (Badreddine et al., 2022; Fino et al., 2018).

##### Electrophysiological recordings

Whole-cell patch-clamp recordings of SPNs were performed with borosilicate glass pipettes (5-8 MΩ) containing (mM): 127 K-gluconate, 13 KCl, 10 HEPES, 10 phosphocreatine, 4 ATP-Mg, 0.3 GTP-Na, 0.3 EGTA (adjusted to pH 7.35 with KOH). Slices were continuously perfused with the extracellular solution containing (mM): 125 NaCl, 2.5 KCl, 25 glucose, 25 NaHCO_3_, 1.25 NaH_2_PO_4_, 2 CaCl_2_, 1 MgCl_2_, 10 μM pyruvic acid bubbled with 95 % O_2_ and 5 % CO_2_. Neurons were visualized under a microscope (Olympus BX51) with a 40x/0.8 water-immersion objective for localizing cells for whole-cell recordings. Signals were amplified using EPC10-2 amplifiers (HEKA Elektronik, Lambrecht, Germany). Current-clamp recordings were filtered at 2.5 kHz and sampled at 5 kHz and voltage-clamp recordings were filtered at 5 kHz and sampled at 10 kHz using the program Patchmaster v2×32 (HEKA Elektronik). Recordings were performed at 32-35 °C to maintain physiological temperature conditions.

##### Optogenetic stimulations

Collimated LEDs placed at the back of the microscope generated wide-field stimulations through a water immersion objective 40X/0.8NA. Activation of ChR2 was performed at λ= 473 nm and consisted of either a single or trains of square pulse of light of various duration (23.2 mW/mm^2^), delivered with a minimum interval of 8 seconds. ChR2 activation led to reliable spiking activity in SOM interneurons (Fino et al., 2018).

### Histology and immunohistochemistry

Mice were deeply anaesthetized with Zoletil/Domitor (2 mL/kg) injected intraperitoneally, then transcardially perfused with first phosphate buffered saline (PBS) and finally 4 % paraformaldehyde (AntigenFix, Diapath). Following perfusion, brains were post-fixed in 4 % paraformaldehyde for 24 hours at 4 °C. Brains were washed with PBS 1X and then kept in PBS at 4°C. Coronal slices (50 µm) obtained with a vibratome (Leica) were used for histology. Brain slices were used for histology and some brain slices for quantification of cFos expressing striatal cells with immunohistochemistry targeting cFos. Slices were blocked with PBST (PBS with 0.3 % Triton X-100) and 5 % (vol/vol) normal goat serum for 1 h 30 and then incubated with the first primary antibody at 4 °C for 24 h (rabbit anti-c-Fos 1:500, Synaptic Systems #226003). The next day, slices underwent three 10 min wash steps in PBS, next they were incubated for 2 h with secondary antibodies (1:200 AlexaFluor 488 anti-rabbit, Invitrogen). Finally, slices underwent three more 10 min wash steps in PBS, followed by mounting and coverslipping with Dako fluorescent mounting medium (Agilent) on microscope slides.

### Drugs

Clozapine-N-oxide (Bio-Techne SAS) was first dissolved in dimethylsulfoxide (DMSO, Sigma, final concentration 25 µg/µL), then aliquoted and stored at −20 °C. For intraperitoneal injections, frozen aliquots were thawed at room temperature on the experimental day, and then further diluted in 0.9 % sterile saline solution to a final concentration of 0.3 µg/µL. The solution was delivered intraperitoneally (3 mg/kg) and, after the injection, the animals were placed back in the home cage for 30 min before the start of the experiment.

## QUANTIFICATION AND STATISTICAL ANALYSIS

### Behavior

#### Accelerating Rotarod

The time to fall (latency) from the accelerating rotarod was recorded to measure the performance of the animals. The latency to fall curves (50 trials) were fitted by a one-phase association model (see (Li and Spitzer, 2020) with the equation Y=Y0 + (Ylim – Y0)*(1-exp(-K*x)) and the values of Y0, Ylim and K (slopes in Fig. 2 and Supplementary Fig. 3) were compared between the tested groups. In addition, a learning index (LI) was computed by subtracting the first 2 trials from the last 2 trials, with LI= Trials_9,10_-Trials_1,2_, in accordance with previous studies (Badreddine et al., 2022; Buitrago et al., 2004; Yin et al., 2009). From the video recordings, several behaviors on the rotarod were measured throughout the training sessions: rearings (animals standing on the rod and lifting its head) and turns (when the animal turns 180° while on the rod). On the videos, we tracked six different key body parts (2 ears, body center, tail base, and 2 limbs) of the mice on the rotarod, using the multiple animal version of DeepLabCut (Lauer et al., 2022). Training was performed using default parameters.

To correct tracking errors we used a custom written code that identified the rotarod barrier on the video frame. We were able in this way to check that each animal was correctly identified (specifically we checked that the tracked bodyparts remain in the same portion of the rotarod). We then analyzed whether scatter plots of the body parts in the XY plane formed well organized clusters. Cluster analysis, used to identify regular behavior, was performed using the standard definition of the silhouette score (Rousseeuw, 1987). This score provides a quantitative measure of how well-defined and distinct the clusters are and can be interpreted as follows: positive values (tending to +1) indicate that data points splits into far apart clusters results whilst a score of zero suggests overlapping clusters or data points equally close to multiple clusters. Silhouettes scores were computed for all mice and for all trials. When it was not possible to compute the score (latency to fall too short to get reliable result), its value was linearly interpolated. Mean silhouette curves obtained for the two groups over the 50 trials of training were then fitted using a one-phase association model (Li and Spitzer, 2020). To assess whether the three parameters of the one-phase association model obtained were different between hM3Dq mice and mCherry mice, we performed a permutation test with 1000 resamplings. Specifically, in each resample we randomly assigned animals to one of the two group, compute the mean silhouette curve, fit the three parameters on the one-phase association model resample data and compute their difference. We then obtained p-values as the proportion of resampled data in which mean differences between the parameters were bigger than true mean differences. All custom written codes used for the clustering analysis and the corresponding relevant data will be posted to online repositories upon publication.

#### Open-field

From the trajectories of the animals in the open field, analysis was performed with Matlab and several parameters extracted. The arena was divided in a 6 x 6 grid with 20 zones at the border and 16 in the center. The position in the grid, the speed, the speed-mode (slow < 8 cm.sec^-1^, fast ≥ 8 cm.sec^-1^) (Granon et al., 2003) and the tortuosity of the mouse trajectories were analyzed at each visit of a zone. Speed is the actual distance ran in a zone divided by the time spent in that zone. The tortuosity is [the actual distance ran in a zone divided by the optimal distance between the points of entry and exit of the animal in that zone] - 1.

### Histology and immunohistochemistry

Images were acquired using an Apotome microscope (Zeiss, Germany). Z-stacks (20-30 µm) with 2 µm step size were acquired with a 20x/0.8 objective. The density of mCherry-expressing or mCherry/cFos co-expressing interneurons was quantified. For each mouse, we acquired Z-stacks on 2-3 different coronal slices in DMS. Using FIJI software, 2 or 3 fields of view per slice (400 x 400 µm) were selected to determine the density of SOM or PV interneurons expressing mCherry only, or co-expressing mCherry and cFos.

### Calcium imaging analysis

GCaMP6f fluorescence signals were analysed with custom-built procedures using R3.5.2 in RStudio environment as previously described and with available codes (Badreddine et al., 2022). The analysis was performed blindly to the training condition and by different experimenters. Briefly, from manually selected ROIs in FIJI software, mean grey values and (x, y) coordinates were extracted for each ROI/slice. Calcium recordings were 700-1000 frames long (around 1 min) and included 7-9 stimulations of cortical afferents. x(t) is the averaged intensity values of pixels in the ROI at time t for one cell. ΔF/F is obtained using y(t) = (x(t) - x0) / x0, where x0 is the mean value of the 50 % lowest values in the last 10 s. ΔF/F was then filtered with a Savitsky-Golay filter of order 3 on sliding windows of 7 frames (0.458 s). For each cell, the amplitude of response to cortical stimulations was calculated by averaging ΔF/F for 5 responses (stimulations #2 to #6). To this aim, fluorescence signals were extracted for each cell on windows of 40 frames (2.6 s) centered on the time of the first maximal amplitudes of ΔF/F (peaks) detected on cells after stimulus. Within this time window, for each response, first the peak of response was detected and then going back to the slope abrupt change between baseline and beginning of the rise, the start was detected. The latency of responses was defined by the timing of the start point of the response, in relation to a T0, given by the first response in the field. The latency was measured for each of the 5 stimulations and the average value was extracted. The amplitude of response was measured as a Delta between the values from the start and the peak points. A cell was defined as active if its amplitude of response was above a threshold defined as M+2SD, with mean (M) and standard deviation (SD) calculated individually for each neuron through the whole recordings; below this threshold, the cell was considered as inactive. The color coded functional maps were extracted with the measure of each cell amplitude within the field of view and inactive cells are represented in white. We distinguished responses from SPNs and other cell types of striatal neurons thanks to a cell-sorting method based on calcium responses we previously developed (Becq G, 2019). To extract the highly active (HA) cells population, we normalized the activity throughout the training conditions by applying a thresholding analysis using the average of the response amplitude in naive animals as reference (calculated from all SPNs in Naïve group). The group containing the cells with the highest amplitude (superior to the threshold) was defined as the HA cells and represents X % of cells in the slice and the others were low-active (LA) cells.

### Electrophysiology

Parameters of electrophysiological intrinsic properties were extracted from the current/voltage curves during the whole-cell recordings. Input resistance was measured by repeated current injections (−20 pA, 500 ms) and rheobase corresponds to the injected current necessary to reach AP threshold. Frequency/current curves were built from AP threshold and averaged frequency was measured for current steps +30 pA above AP threshold. SPNs were held at their physiological membrane potential and there was no statistical difference in the holding membrane potentials between the different experimental conditions. For optogenetic experiments, light-induced current were recorded at −40mV, a potential above chloride reversal (set at −60mV, the SPN physiological chloride reversal). The amplitude of light-induced currents was measured as a delta between the start to the peak of the outward current triggered by a 5-10 ms LED pulse. To quantify the modulation of SPN frequency discharge by SOM cells, a suprathreshold 1 sec long depolarizing step induced firing in SPNs and the LED was ON for 300ms at the center of the depolarizing step. The frequency ratio correspond to the spike frequency during or after LED divided by the control frequency before LED. Data analysis was carried out in Fitmaster (HEKA Elektronik, Germany).

### Statistical analysis

The data is presented and plotted as mean ± SEM (unless otherwise stated), where SEM refers to standard error of the mean. p values are represented by symbols using the following code: * for p< 0.05, ** for p< 0.01, *** for p< 0.001. Exact p values and statistical tests are stated in the figure legends or in the core of the manuscript. Statistical analysis was performed using Prism 5.0 (GraphPad, San Diego, USA), R or Python. The sample size for the different sets of data is mentioned in the text or in the respective figure legends. Normality of each data set was checked using D’Agostino and Pearson’s test. Statistical significance was assessed using Student’s t-test or Mann-Whitney’s U-test and Wilcoxon’s signed rank test for unpaired and paired data, respectively. One-way Anova was used to compare all the effects in DMS between the different training conditions. Pearson correlation was used for relationship between percentage of HA cells and learning index. Two-way Anova followed by Bonferroni *post-hoc* correction was used to compare different parameters (calcium dynamics) evolving throughout the training conditions and learning curves in different treatment conditions.

## ADDITIONAL SECTION

## Author contribution

SR, GZ and EF designed experiments. SR, GZ, NB, FA and EF performed experiments. SR, GZ, FA, NB, SS, IB and EF performed analysis of behavioral (SR, GZ, NB, SS and IB), calcium imaging (GZ, FA and EF) and electrophysiological (EF) data. SR, GZ, NB and EF wrote the manuscript; all authors read and edited the manuscript. EF supervised the project and acquired funding.

## SUPPLEMENTARY INFORMATION

**Supplementary Fig. 1, related to Figure1.**
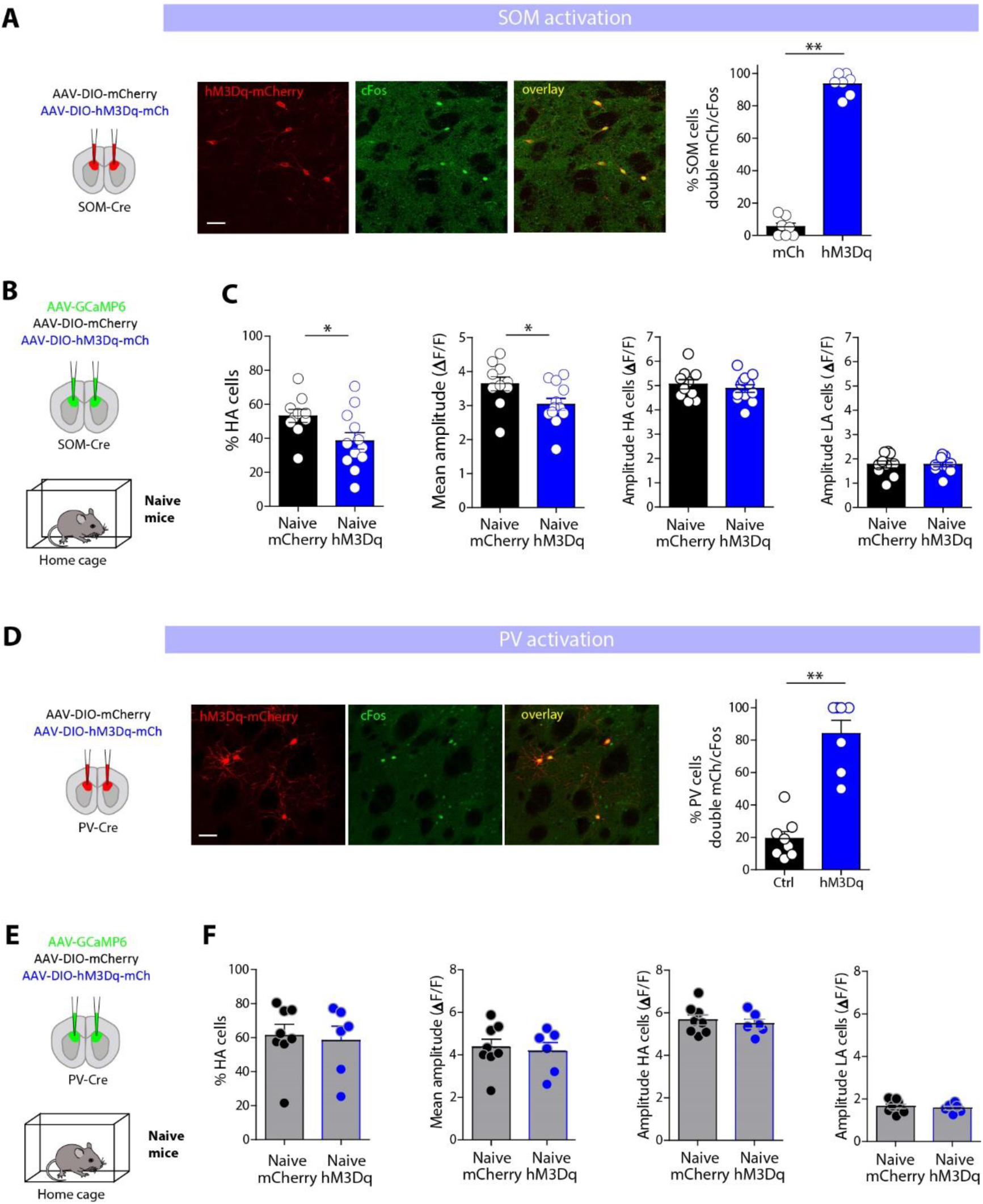
A, After injections in SOM-cre mice of AAV-DIO-mCherry or AAV-DIO hM3Dq-mCherry, SOM cells express either constructs. Immunostainings of mCherry, cFos, and overlay in SOM cells (scale bar: 20µm). After CNO injections only SOM cells (expressing hM3Dq) are activated as double expressing cFos and mCherry in SOM cells is around 95% in hM3Dq animals and only 5% in mCherry mice (p=0.0021, Mann-Whitney). B, Experimental conditions. SOM-cre mice are injected with GCaMP6f and either floxed mCherry (control group, n=6 mice) or floxed hM3Dq (SOM-activated group, n=7 mice). Both groups of naïve animals received CNO injection and put back in their home cage. The day after, *ex vivo* calcium imaging was performed on control naïve mCherry group (mCherry, black, n=10 slices from 6 mice) or naïve with SOM activated (hM3Dq, blue, n= 12 slices from 7 mice). C, Mean amplitudes of all SPNs responses in the recording field, percentage of HA cells, amplitude of HA and LA cells for both groups. There is a significant decrease of mean amplitude (p= 0.0378, Mann-Whitney) and percentage of HA cells (p= 0.0443) between both naïve groups, with no change in the amplitude of HA (p= 0.6209) or LA (p= 0.7169) cells. Comparing to the conditions of early-training induced changes in amplitude and percentage of HA cells (shown in Fig.1), both the decrease in the percentage of HA cells (p= 0.0141) and amplitude (p=0.0226) are lower in naïve SOM-stimulated mice compared to early-trained mice, indicating that SOM inhibition needs to be coupled to training to form representations of early learning. D, After injections in PV-cre mice of AAV-DIO-mCherry or AAV-DIO hM3Dq-mCherry, PV cells express either constructs. Immunostainings of mCherry, cFos, and overlay in PV cells (scale bar: 20µm). After CNO injections, PV cells (expressing hM3Dq) are mainly activated as double expressing cFos and mCherry in PV cells is around 80% in hM3Dq animals and only 20% in mCherry mice (p= 0.0003, Mann-Whitney). E, Experimental conditions. PV-cre mice are injected with GCaMP6f and either floxed mCherry (control group) or floxed hM3Dq (PV-activated group). Naïve animals receive CNO injection and put back in their home cage. The day after, *ex vivo* calcium imaging is performed on control naïve group (mCherry, black) or naïve with PV activated (hM3Dq, blue). F, Mean amplitudes of all SPNs responses in the recording field, percentage of HA cells, amplitude of HA and LA cells for both groups. There is no significant difference in the mean amplitude (p= 0.8518, Mann-Whitney), percentage of HA cells (p= 1.000), amplitude of HA (p= 0.7546) or LA (p= 0.7546) cells between both naïve groups.

**Supplementary Fig.2, related to Figure2.**
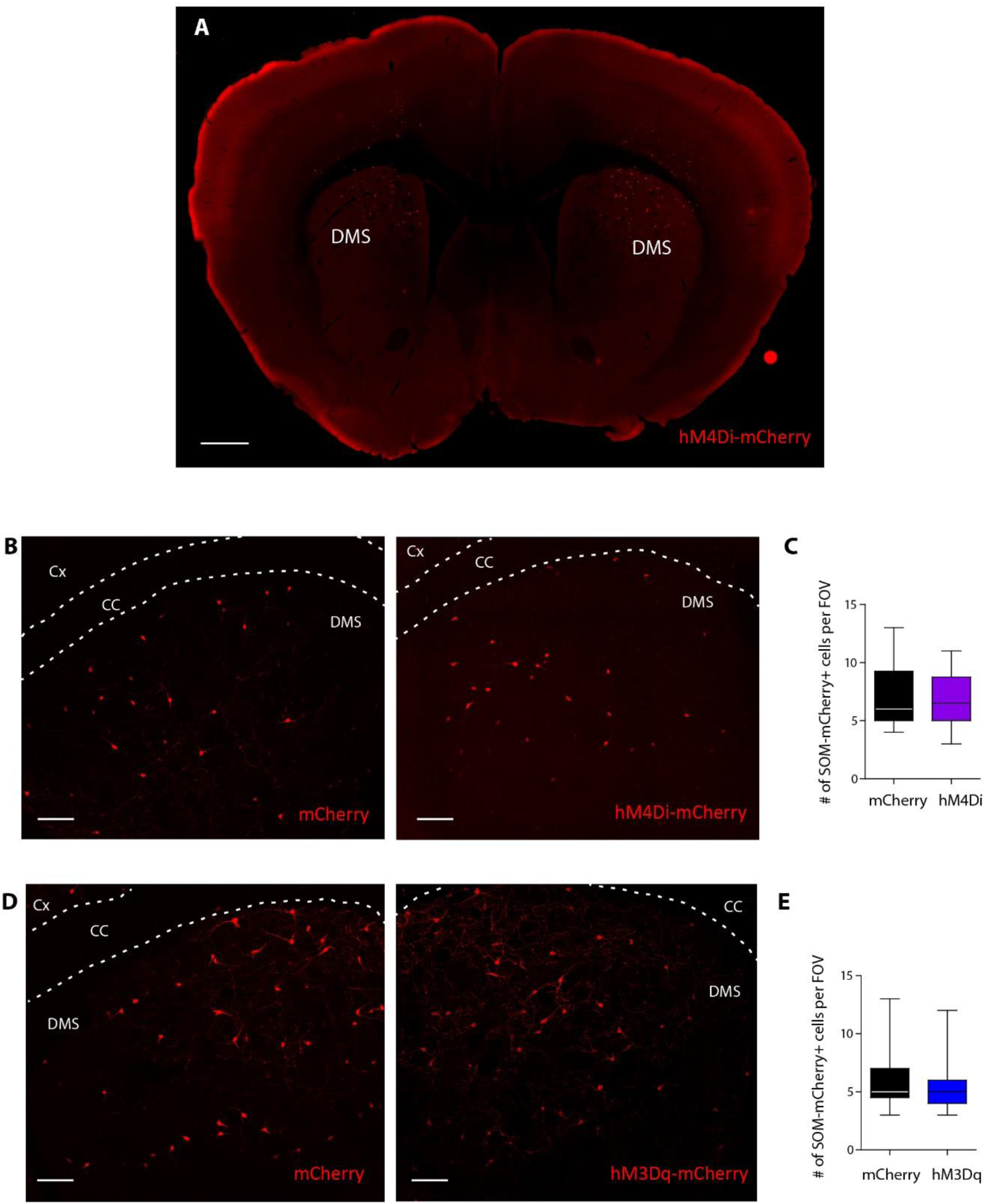
A, Wide field image of hM4Di-mCherry expression within DMS after bilateral injections. Scale bars: 500 µm. B, Confocal images of mCherry-expressing cells or hM4Di-mCherry expressing cells in SOM cells. Scale bars: 100 µm. C, Quantification of mCherry+ and hM4Di-mCherry-expressing cells: the number of cells is expressed per field of view (FOV) of 400 x 400 µm and the quantification of fluorescent cells was done in each mouse in 1-2 slices and 2-3 FOV. There was no significant difference in the density of mCherry+ cells (black, 7±0.6 cells, n=18 FOV, n=6 mice) and hM4Di-mCherry+ cells (purple, 6.9±0.5 cells, n=28 FOV, n=9 mice) (p=0.9114, t-test). D, Confocal images of mCherry-expressing cells or hM3Dq-mCherry expressing cells in SOM cells. Scale bars: 100 µm. E, Quantification of mCherry+ and hM3Dq-mCherry-expressing cells: the number of cells is expressed per field of view (FOV) of 400 x 400 µm and the quantification of fluorescent cells was done in each mouse in 1-2 slices and 2-3 FOV. There was no significant difference in the density of mCherry+ cells (black, 5.9±0.5 cells, n=25 FOV, n=7 mice) and hM3Dq-mCherry+ cells (blue, 5.5±0.4 cells, n=28 FOV, n=7 mice) (p=4604, t-test).

**Supplementary Figure 3, related to Figure 2.**
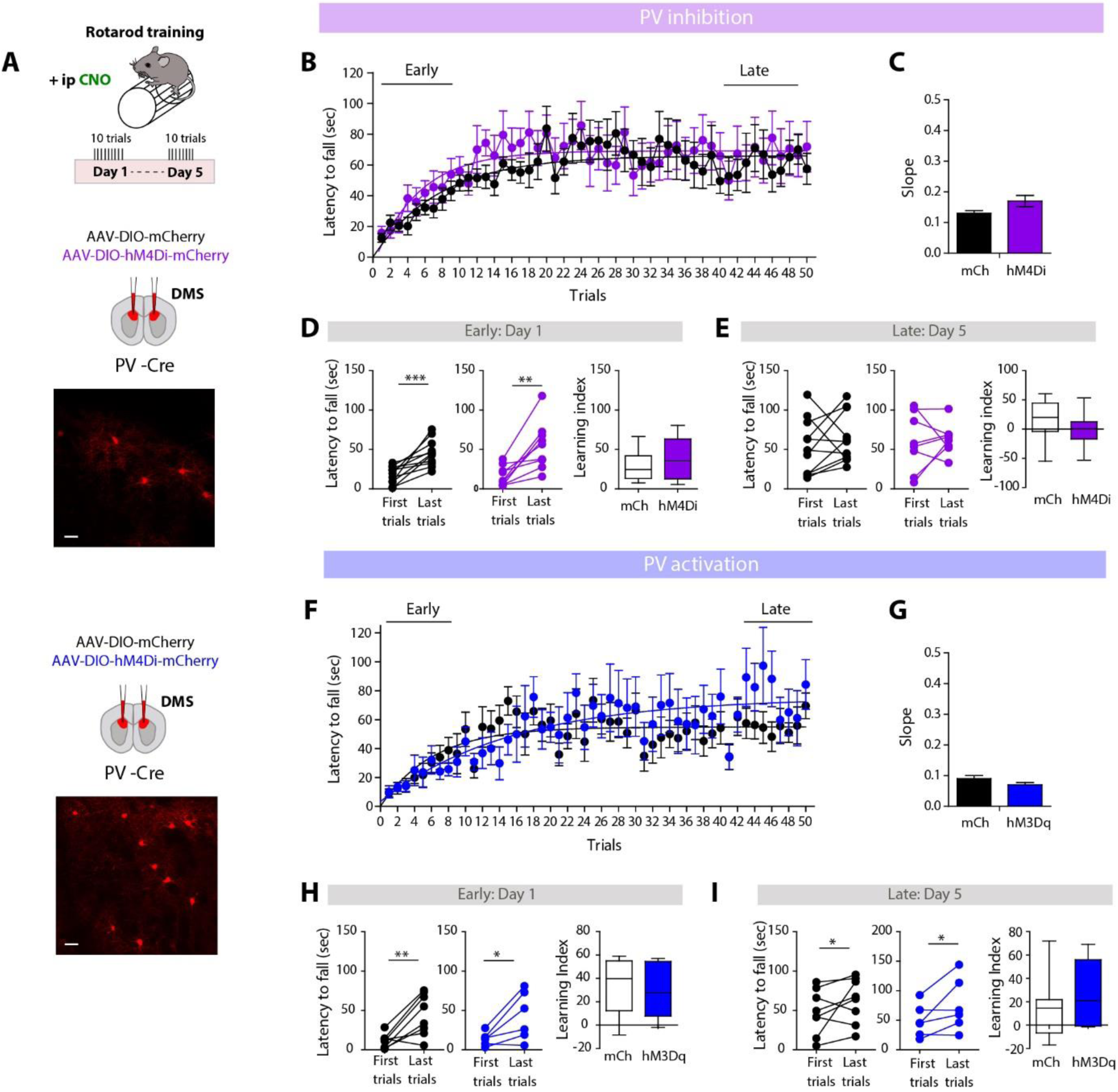
A, Experimental conditions, PV-cre mice are injected with either floxed mCherry (control group) or floxed hM4Di (PV silencing group) or floxed hM3Dq (PV activating group) during rotarod training (5 days, 1 session with 10 trials per day). Confocal images of PV cells expressing hM4Di-mCherry or hM3Dq-mCherry (scale bar: 20µm). B, Learning curves and one-phase association fits of the control mice (mCherry, black, n=11 mice) or PV silenced mice (hM4Di, purple, n=8 mice) throughout the 5 training sessions (50 trials). C, Slopes of the one-phase association fits for both groups are not significantly different (p= 0.0766). D, In early (Day 1) training, significant improvement in the mice performance between the first and last trials for both groups and similar learning index (p=0.4703). E, Both groups reach similar performance in the late phase of the training (Day 5) with similar learning index (p=0.3893). F, Learning curves and one-phase association fits of the control mice (mCherry, black, n= 6 mice) or PV activated mice (hM3Dq, blue, n= 6 mice) throughout the 5 training sessions (50 trials). G, Slopes of the one-phase association fits for both groups (p= 0.1390). H, In early (Day 1) training, significant improvement in the mice performance between the first and last trials for both groups with similar learning index (p=0.8518). I, Both groups then reach similar performance by the late phase of the training (Day 5) with similar learning index (p= 0.5728).

**Supplementary Figure 4, related to Figure 2.**
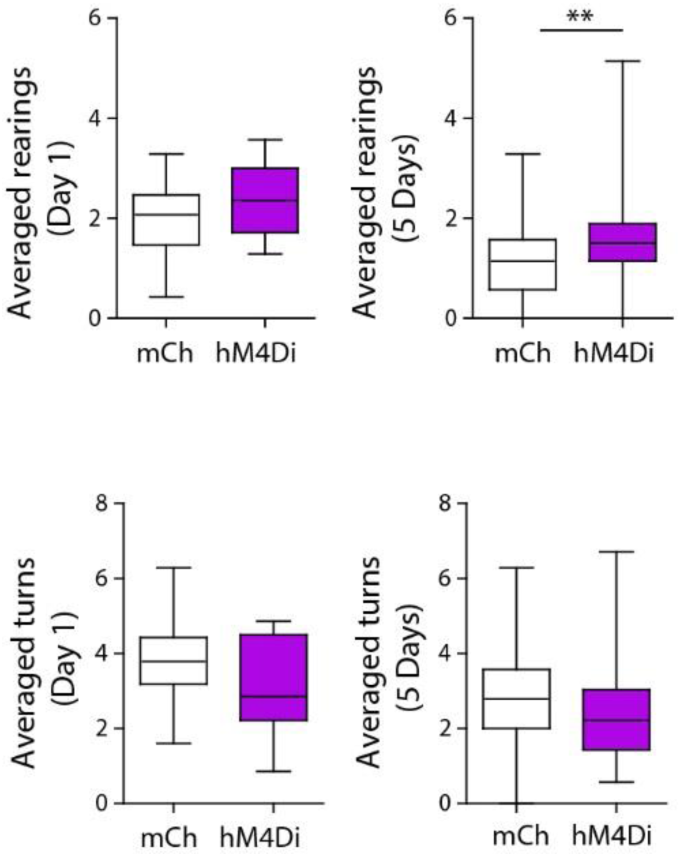
Numbers of rearings and turns (180° turns) the mice displayed during the rotarod training. Over the overall training sessions (5 days), hM4Di mice (n=8 mice) had significantly (p= 0.003, Mann-Whitney) more rearings than the control group (n=7 mice). The number of turns was not significantly different.

**Supplementary Figure 5, related to Figure 3.**
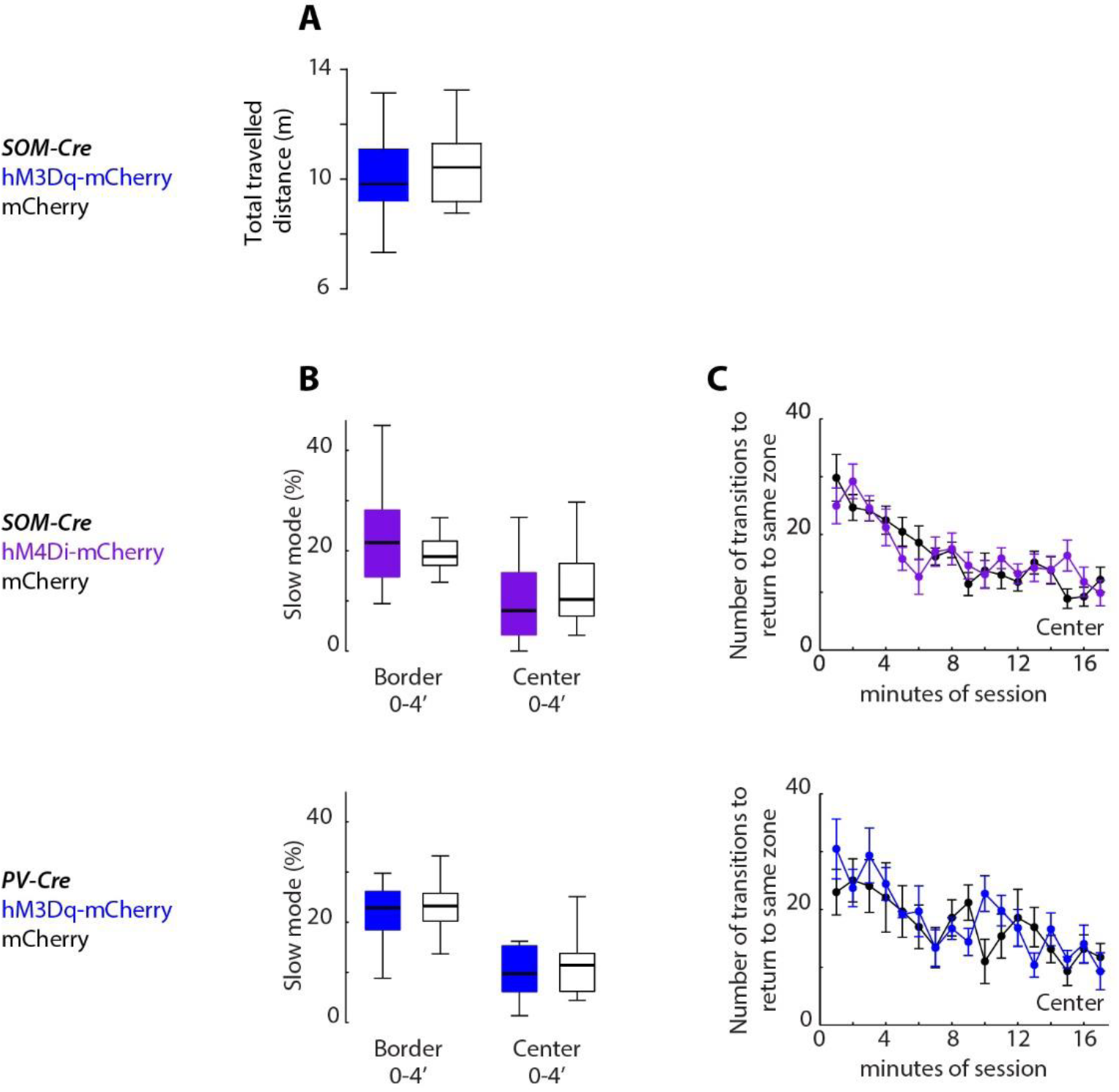
A, No statistical difference in the total travelled distance between hM3Dq (n=11) and mCherry (n=13) mice. B-C, fraction of zones visited in slow-mode during the first 4 minutes (B) and number of transitions before returning to a same zone of the arena (C) for SOM-Cre mice (top) injected with AAV-DIO-hM4Di-mCherry (purple, n=17 mice) or AAV-DIO-mCherry (black, n=12 mice) and for PV-Cre mice (bottom) injected with AAV-DIO-hM3Dq-mCherry (blue, n=8 mice) or AAV-DIO-mCherry (black, n=8 mice).

